# CAPZ, but not canonical autophagy, regulates the endosomal-exosomal trafficking of plasma membrane PD-L1

**DOI:** 10.64898/2026.03.12.711228

**Authors:** Peng Xu, Zuodong Ye, Yanni Zhu, Naixin Lin, Chuenfuk Chan, Chengyi Zhu, Xiaoyin Zhang, Yijing Wang, Wei Sun, Meiyu Peng, Yingying Lu, Jianbo Yue

**Author notes:** equal contribution (Peng Xu, Zuodong Ye). Corresponding author: Jianbo Yue, Yingying Lu.

## Abstract

Programmed death-ligand 1 (PD-L1) undergoes continuous endocytosis and intracellular trafficking that regulate its degradation, recycling, and exosomal release. Although PD-L1 internalization is known to depend on RAB5-mediated endocytosis, whether canonical autophagy contributes to its subsequent endosomal trafficking remains unclear. Here we show that the endosomal and exosomal trafficking of cell surface PD-L1 occurs independently of the canonical autophagy machinery. Genetic disruption of core autophagy components, including LC3B, ATG4B, ATG5, and ATG7, did not impair trafficking of internalized PD-L1 to early endosomes, multivesicular bodies, late endosomes, or extracellular vesicles. Pharmacologic inhibition of autophagosome–lysosome fusion enhanced accumulation of PD-L1 within RAB5-and CD63-positive compartments, but this effect persisted in cells lacking LC3B, ATG5, or ATG7, indicating that PD-L1 trafficking through the endosomal–exosomal pathway does not require canonical autophagy. Instead, we identify the actin-capping protein CAPZ as a key regulator of an endosomal maturation checkpoint that controls PD-L1 sorting. Loss of CAPZ impaired progression of PD-L1 from early to late endosomal compartments, reduced PD-L1 incorporation into multivesicular bodies and extracellular vesicles, and redirected PD-L1 toward RAB11-dependent recycling, resulting in increased plasma membrane PD-L1 abundance. These findings establish that PD-L1 membrane fate is determined by CAPZ-dependent endosomal maturation rather than canonical autophagy and identify endosomal trafficking machinery as a critical regulator of immune checkpoint distribution.

## INTRODUCTION

The programmed death-1 (PD-1) receptor and its ligand PD-L1 constitute a central immune checkpoint pathway that restrains T cell activation and maintains peripheral tolerance^1,2^. Engagement of PD-L1 on tumor or antigen-presenting cells with PD-1 on activated T cells attenuates TCR signaling, suppresses cytokine production, and promotes T cell exhaustion, thereby enabling immune evasion in cancer^3^. Therapeutic blockade of the PD-1/PD-L1 axis has transformed oncology, producing durable responses across multiple tumor types^4–6^. However, clinical responses remain heterogeneous, and both intrinsic and acquired resistance are common^7^. Beyond transcriptional control of PD-L1 expression, emerging evidence indicates that post-translational regulation and membrane trafficking critically determine PD-L1 surface abundance and, consequently, checkpoint potency^8^.

PD-L1 expression is regulated at multiple levels. Oncogenic signaling pathways, including PI3K–AKT–mTOR and MAPK cascades, enhance PD-L1 transcription and stability^9,10^, while inflammatory cytokines such as IFN-γ drive inducible expression via JAK–STAT signaling^11,12^. Post-translational modifications, including glycosylation, ubiquitination, and palmitoylation, further modulate PD-L1 stability and turnover^13–16^. Importantly, PD-L1 is not statically displayed at the plasma membrane but undergoes constitutive internalization and recycling. Endocytic trafficking determines whether PD-L1 is recycled back to the cell surface, delivered to lysosomes for degradation, or sorted into multivesicular bodies (MVBs) for exosomal secretion. Thus, endosomal sorting decisions directly control immune checkpoint availability^17–20^.

Autophagy, a conserved lysosome-dependent degradation pathway, has been implicated in immune regulation and tumor progression^21,22^. Canonical macroautophagy involves formation of double-membraned autophagosomes through coordinated action of ATG proteins, including ATG5, ATG7, ATG4, and LC3 family members, followed by autophagosome–lysosome fusion^23^. Several studies have suggested that autophagy influences PD-L1 levels, either by promoting its degradation or by altering its intracellular distribution under stress conditions^24^. Pharmacologic or genetic modulation of autophagy components has been reported to affect PD-L1 abundance in certain contexts, leading to the proposal that autophagy machinery may participate in PD-L1 turnover^19,24–26^. However, whether canonical autophagy directly governs the endosomal–exosomal trafficking itinerary of plasma membrane PD-L1 remains unclear. In particular, it is unknown whether core autophagy proteins function as essential determinants of PD-L1 progression through the endosomal maturation cascade.

Endosomal maturation is orchestrated by sequential Rab GTPase transitions and coordinated membrane remodeling events that regulate cargo sorting. Early endosomes marked by RAB5 mature into late endosomes and MVBs through Rab conversion and acquisition of late endosomal markers such as LAMP1 and CD63. Cargo can be recycled to the plasma membrane via RAB11-positive recycling endosomes, targeted for lysosomal degradation, or incorporated into intraluminal vesicles destined for exosomal release^27–29^. The molecular checkpoints that determine PD-L1 sorting at these maturation stages have not been fully defined^30,31^.

CAPZ (F-actin–capping protein) is a conserved heterodimer composed of α and β subunits that binds to the barbed ends of actin filaments to regulate actin polymerization dynamics^32,33^. Through its canonical role in actin capping, CAPZ influences cytoskeletal organization, cell migration, and membrane remodeling^34–36^. However, accumulating evidence indicates that CAPZ functions beyond cytoskeletal regulation^37–39^. In addition to modulating cortical actin architecture, CAPZ has been implicated in membrane trafficking processes. Our previous work demonstrated that CAPZ directly participates in endosomal maturation and regulates the Rab5–Rab7 transition, thereby controlling cargo progression through the endolysosomal pathway. These findings suggest that CAPZ may act as a regulatory component of the endosomal machinery independent of its classical structural role^40,41^.

Here, we examined whether PD-L1 trafficking is governed by canonical autophagy or by endosomal maturation mechanisms. We found that core autophagy machinery is dispensable for PD-L1 endosomal–exosomal transport. Instead, CAPZ functions as a critical regulator of an endosomal maturation checkpoint that determines whether PD-L1 is recycled, degraded, or secreted. These findings redefine PD-L1 regulation as a process controlled by endosomal sorting rather than autophagy.

## RESULTS

### The endosomal trafficking of cell surface PD-L1 is independent of the canonical autophagy machinery

To determine whether canonical autophagy contributes to the intracellular trafficking of cell surface PD-L1, we first defined the endosomal itinerary of internalized PD-L1. Using the antibody-feeding–based surface labeling assay previously described^17^, surface PD-L1 was labeled at 4°C and internalization was initiated by shifting cells to 37°C. Internalized PD-L1 rapidly accumulated in EEA1-positive early endosomes at 30 min (**Fig. S1A**), followed by redistribution at later time points, consistent with endosomal maturation. In parallel, PD-L1 progressively colocalized with LAMP1-positive late endosomal/lysosomal compartments (**Fig. S1B**). Quantitative analyses confirmed time-dependent increases in colocalization coefficients with both early and late endosomal markers. These data establish that cell surface PD-L1 undergoes canonical endosomal trafficking from early to late compartments following internalization.

Given this endolysosomal progression, we next asked whether LC3-positive vesicles participate in PD-L1 trafficking. Minimal PD-L1–LC3 overlap was observed at baseline; however, significant colocalization emerged at 60 and 120 min after cell surface PD-L1 internalization (**Fig. 1A**), suggesting that PD-L1 dynamically associates with LC3-positive compartments during intracellular trafficking. To further evaluate this association, we examined the impact of autophagic flux inhibition. Under basal conditions, PD-L1 partially colocalized with LC3B puncta (**Fig. S1C**, upper panel). Treatment with Bafilomycin A1, a vacuolar H⁺- ATPase inhibitor that blocks lysosomal acidification and prevents fusion of lysosomes with autophagosomes or late endosomes^42^, markedly increased LC3B puncta formation and strongly enhanced PD-L1–LC3B colocalization, consistent with accumulation of LC3-positive autophagic structures. These findings suggested that autophagy might participate in PD-L1 trafficking. To confirm that this signal reflected bona fide LC3-positive membranes, we depleted ATG4B, a protease required for LC3 maturation^43^. ATG4B knockdown abolished LC3B puncta formation and eliminated PD-L1–LC3B colocalization (**Fig. S1**C, lower panel). Efficient ATG4B depletion and disruption of autophagic flux were verified by immunoblotting (**Fig. S1D–S1E**). These data indicate that PD-L1 dynamically associates with LC3-positive vesicles in an LC3 processing– dependent manner, raising the possibility that canonical autophagy contributes to PD-L1 endosomal trafficking.

**Figure 1.**
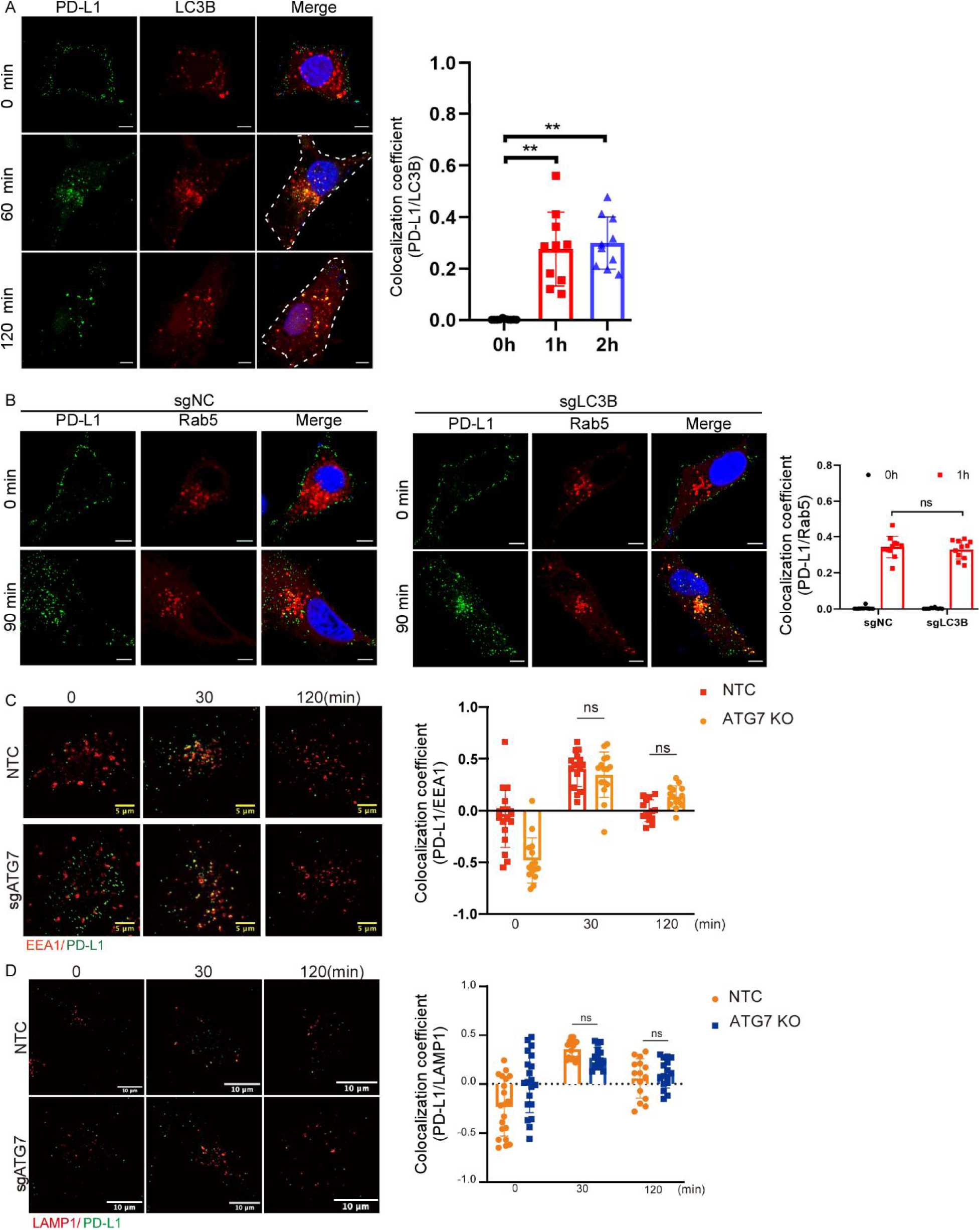
Internalized PD-L1 associates with LC3-positive vesicles but traffics independently of canonical autophagy. (A) HeLa cells were subjected to antibody-feeding assays to label cell surface PD-L1 at 4°C followed by internalization at 37°C for the indicated times. Immunofluorescence analysis shows colocalization of internalized PD-L1 with LC3B. Quantification of Manders’ coefficients is shown at right. (B) Colocalization of internalized PD-L1 with RAB5-positive early endosomes in control or LC3B knockdown cells. Quantification of colocalization coefficients is shown. (C–D) Colocalization of internalized PD-L1 with EEA1 (C) and LAMP1 (D) in control and ATG7-deficient cells at indicated time points. Quantification of colocalization coefficients is shown. The bar graphs in panels (A-D) show data that represent the mean ± s.e.m of three independent experiments, and the asterisks indicate significant differences at ** P < 0.01. ‘ns’ indicates that data were not significantly different.

To directly test whether LC3 functionally regulates PD-L1 trafficking, we depleted LC3B and examined PD-L1 localization relative to Rab5-positive early endosomes. LC3B depletion was confirmed by immunoblotting, with expected alterations in autophagic flux markers (**Fig. S1F**). Despite effective LC3B reduction, internalized PD-L1 displayed normal colocalization with Rab5 at 90 min (**Fig. 1B**), and quantitative analysis revealed no significant difference compared with control cells. We then disrupted ATG7, an essential enzyme required for LC3 lipidation and autophagosome formation^44^. Loss of ATG7 and impairment of LC3 processing were confirmed by immunoblotting (**Fig. S1G**). Notably, ATG7 deficiency did not alter PD-L1 trafficking kinetics: internalized PD-L1 colocalized with EEA1-positive early endosomes and subsequently with LAMP1-positive late compartments with dynamics comparable to control cells (**Fig. 1C–1D**). These data indicate that, although PD-L1 transiently associates with LC3-positive vesicles, canonical autophagy components are not required for its progression through the endosomal system.

Having established that canonical autophagy is dispensable for basal PD-L1 trafficking, we next asked whether acute pharmacological blockade of autophagic flux alters PD-L1 trafficking dynamics. We utilized the triazine compound 6J1, previously characterized as an inhibitor of autophagosome–lysosome fusion^45,46^. Immunoblot analysis revealed dose-dependent accumulation of LC3-II and p62 in HeLa cells treated with 6J1 (**Fig. S2A**), consistent with inhibition of autophagic flux. In RFP-GFP-LC3 reporter cells, 6J1 markedly increased yellow puncta without accumulation of red-only puncta (**Fig. S2B**), confirming impaired autophagosome maturation and fusion with lysosomes. These effects phenocopied Bafilomycin A1 and established 6J1 as a potent autophagy flux inhibitor.

Strikingly, 6J1 markedly increased the colocalization of PD-L1 with LC3-positive puncta, as shown by both PD-L1-GFP/LC3-RFP coexpression and immunostaining of endogenous PD-L1 and LC3B (**Figs. 2A** and left panel in **2B**). To determine whether this colocalization reflected authentic LC3-positive structures, we disrupted ATG5, a core component required for LC3 lipidation^47^. ATG5 knockdown efficiently reduced ATG5 expression (**Fig. S2C**) and impaired autophagic flux, as evidenced by defective LC3 processing and altered p62 accumulation (**Fig. S2D**). Under these conditions, 6J1-induced PD-L1–LC3B colocalization was largely lost (right panel in **Fig. 2B**), confirming the specificity of the LC3 signal associated with PD-L1-containing puncta. In contrast, 6J1-induced colocalization of PD-L1 with Rab5-positive early endosomes was preserved in LC3B-deficient cells (**Fig. 2C**). Consistent with this observation, 6J1 also increased PD-L1 colocalization with EEA1 in both control and ATG7-deficient cells (**Fig. 2D**). These findings indicate that, although 6J1 promotes the accumulation of PD-L1 in LC3-positive compartments, trafficking of internalized PD-L1 to early endosomes does not require the canonical autophagy machinery.

**Figure 2.**
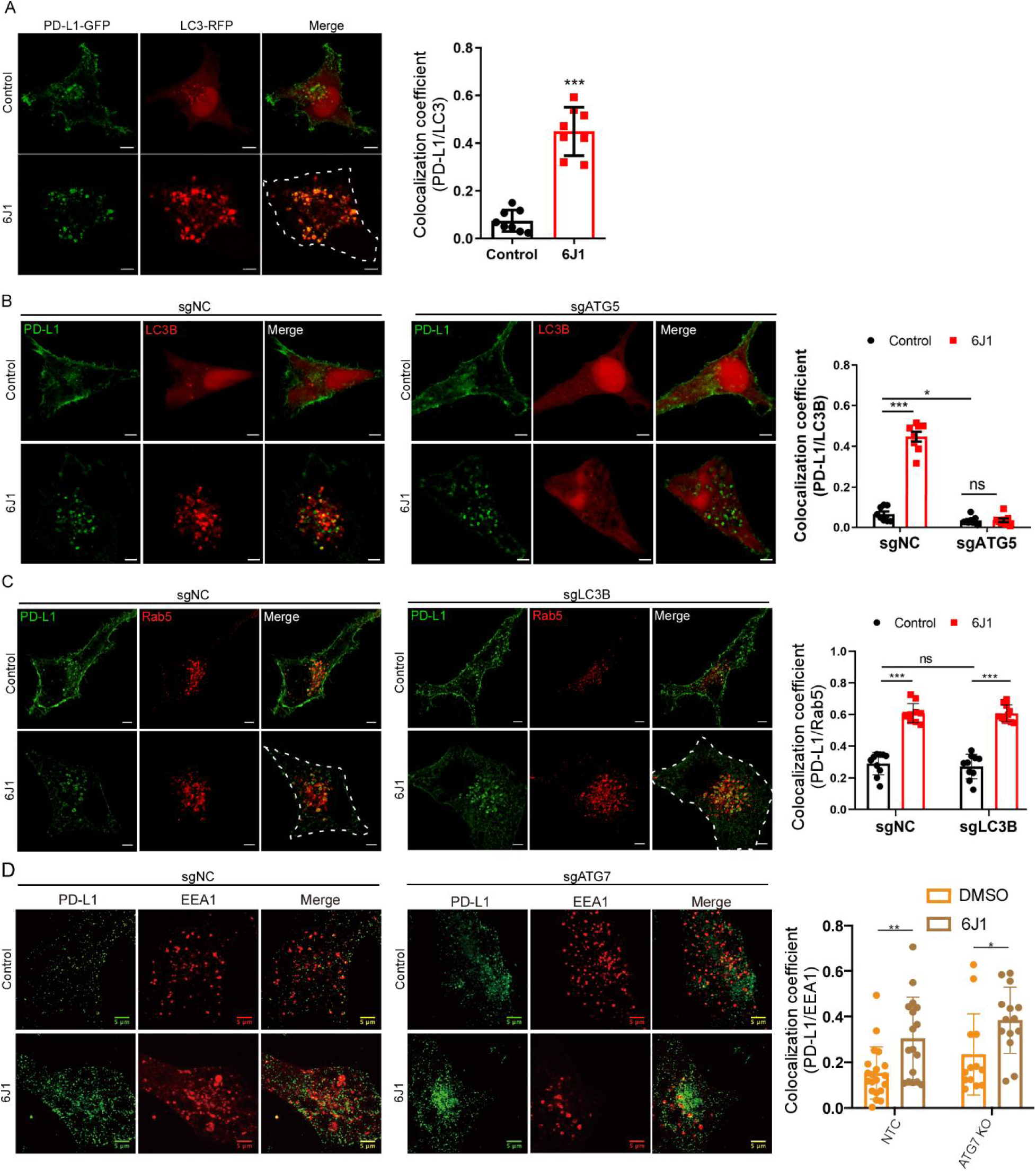
6J1 promotes PD-L1 accumulation in LC3-positive structures, whereas trafficking of PD-L1 to early endosomes is autophagy-independent. **(A**) HeLa cells coexpressing PD-L1-GFP and LC3-RFP were treated with DMSO or 6J1 (1 mM) and analyzed by confocal microscopy. Scale bar, 10 μm. (**B**) Control and ATG5-deficient HeLa cells were treated with DMSO or 6J1 (1 mM), followed by immunostaining for endogenous PD-L1 and LC3B. Scale bar, 10 μm. (**C**) Control and LC3B-deficient HeLa cells were treated with DMSO or 6J1 (1 mM), followed by immunostaining for PD-L1 and Rab5. Scale bar, 10 μm. (**D**) Control and ATG7-deficient HeLa cells were treated with DMSO or 6J1 (1 mM), followed by immunostaining for PD-L1 and EEA1. Scale bar, 5 μm. Data are presented as mean ± SEM. Statistical significance was determined as indicated in the figure. ns, not significant; *p < 0.05; **p < 0.01; ***p < 0.001.

Together, these results indicate that although inhibition of autophagic flux enhances the association of PD-L1 with LC3-positive structures, trafficking of internalized PD-L1 to early and late endosomal compartments does not require the canonical autophagy machinery.

### The exosomal trafficking of cell surface PD-L1 is independent of canonical autophagy machinery

Having demonstrated that canonical autophagy is dispensable for endosomal trafficking of internalized PD-L1 (**Figs. 1** and **2**), we asked whether the autophagy machinery contributes to the exosomal trafficking of membrane PD-L1. PD-L1 has been reported to be secreted via extracellular vesicles (EVs), including exosomes^48,49^. Consistent with this, PD-L1 was readily detected in EVs isolated from both HeLa and 4T1 cells (**Figs. S3A** and **S3B**), together with established exosomal markers TSG101, ALIX, CD63, and CD9, whereas the endoplasmic reticulum marker calnexin was absent, confirming the purity of the preparations.

To determine whether internalized membrane PD-L1 traffics through multivesicular bodies (MVBs), we examined its localization relative to CD63, a canonical MVB/exosomal marker^50^. Endocytosed PD-L1 exhibited strong colocalization with CD63 at later time points (left panel of **Figs. 3A and 3B**), indicating delivery of membrane PD-L1 to MVB compartments. In parallel, PD-L1 robustly colocalized with RAB27A (left panel of **Fig. S3C**), a Rab GTPase required for exosome secretion^51^, further supporting engagement of the exosomal secretion machinery. Because amphisomes arise from fusion of autophagosomes with late endosomes/MVBs and have been implicated in EV release^52^, we next tested whether canonical autophagy participates in exosomal PD-L1 trafficking. Depletion of LC3B did not alter PD-L1 colocalization with CD63 (right panel of Fig. 3A) or with RAB27A (right panel of **Fig. S3C**), and quantitative analyses revealed comparable colocalization coefficients between control and LC3B-deficient cells. Likewise, depletion of ATG5 or ATG7 did not affect PD-L1 colocalization with CD63 (bottom panel of **Fig. 3A** and right panel of **Fig. 3B**, respectively), indicating that LC3 lipidation and autophagosome formation are not required for PD-L1 delivery to MVBs.

**Figure 3.**
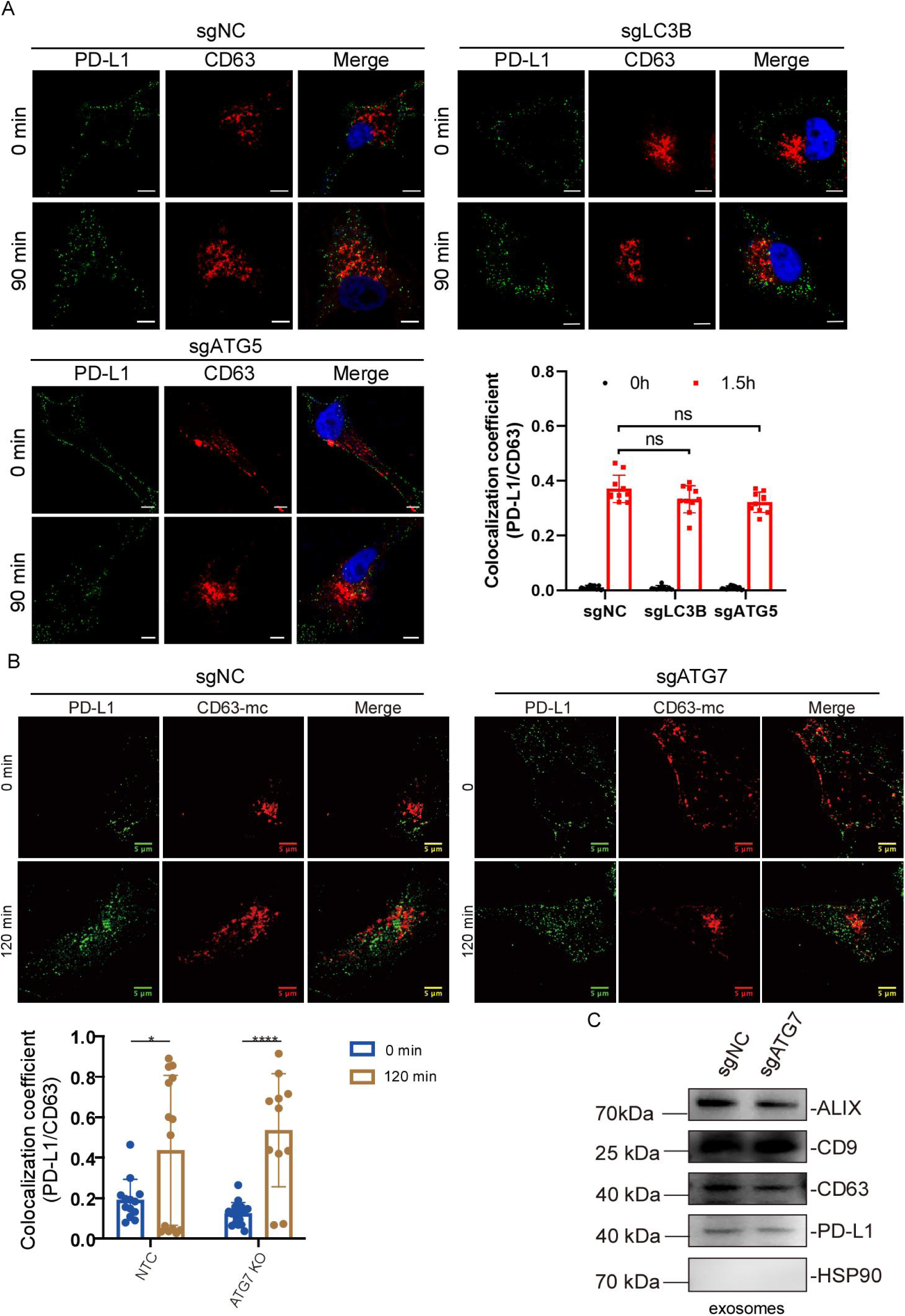
Canonical autophagy is dispensable for trafficking of internalized PD-L1 to CD63-positive multivesicular bodies and for its incorporation into exosomes. (A) Control, LC3B-deficient, and ATG5-deficient HeLa cells were incubated with anti-PD-L1 antibody on ice to label cell surface PD-L1 and then chased at 37 °C for the indicated times. Cells were fixed and immunostained for PD-L1 and CD63. Representative confocal images and quantification of internalized PD-L1 increasingly colocalized with the MVB/exosomal marker CD63 are shown. Scale bars, 10 μm. (B) Control and ATG7-deficient HeLa cells were labeled for cell surface PD-L1 and chased for the indicated times, followed by immunostaining for PD-L1 and CD63. Representative images and quantification of PD-L1 colocalization with CD63 are shown. Scale bars, 5 μm. (C) Exosomes were isolated from conditioned medium of control and ATG7-deficient HeLa cells and analyzed by immunoblotting for PD-L1 and the exosomal markers ALIX, CD9, and CD63. HSP90 was used as a negative control. Data are presented as mean ± SEM. Statistical significance was analyzed as indicated in the figure. ns, not significant; *p < 0.05; ****p < 0.0001.

We next asked whether canonical autophagy is required for exosomal secretion of PD-L1. Exosomes isolated from control and ATG7-deficient cells contained comparable levels of PD-L1, and the abundance of the exosomal markers ALIX, CD9, and CD63 was unchanged (**Fig. 3C**), indicating that loss of canonical autophagy does not impair incorporation of PD-L1 into exosomes. We further examined whether acute stimulation of EV release affects PD-L1 abundance and whether this response depends on autophagy. Ionomycin treatment markedly reduced total cellular PD-L1 levels, consistent with Ca²⁺-stimulated vesicular secretion^53^. Importantly, this ionomycin-induced reduction was not prevented by depletion of ATG4B or ATG7 (**Figs. S3D–S3E**), indicating that Ca²⁺-triggered PD-L1 loss occurs independently of canonical autophagy.

Collectively, these findings further indicate that canonical autophagy is not required for trafficking of PD-L1 to MVBs or for its secretion via exosomes.

### The endosomal trafficking of cell surface PD-L1 is dependent on the CAPZ

Having established that canonical autophagy is dispensable for PD-L1 endosomal and exosomal trafficking (**Figs. 1–3**), we next sought to identify the molecular machinery that functionally regulates PD-L1 endosomal sorting. Given the role of CAPZ in endosomal dynamics^40,41,54^, we examined whether CAPZ participates in PD-L1 trafficking. Internalized PD-L1 dynamically associated with CAPZβ-positive structures. At 30 min following internalization, PD-L1 puncta exhibited marked overlap with CAPZβ, and quantitative analysis revealed a significant increase in Manders’ coefficients relative to baseline (**Fig. 4A**). In addition, 6J1 treatment markedly enhanced the colocalization of PD-L1 with CAPZβ (**Fig. 4B**), further supporting the idea that CAPZ is closely associated with PD-L1-containing endosomal structures.

**Figure 4.**
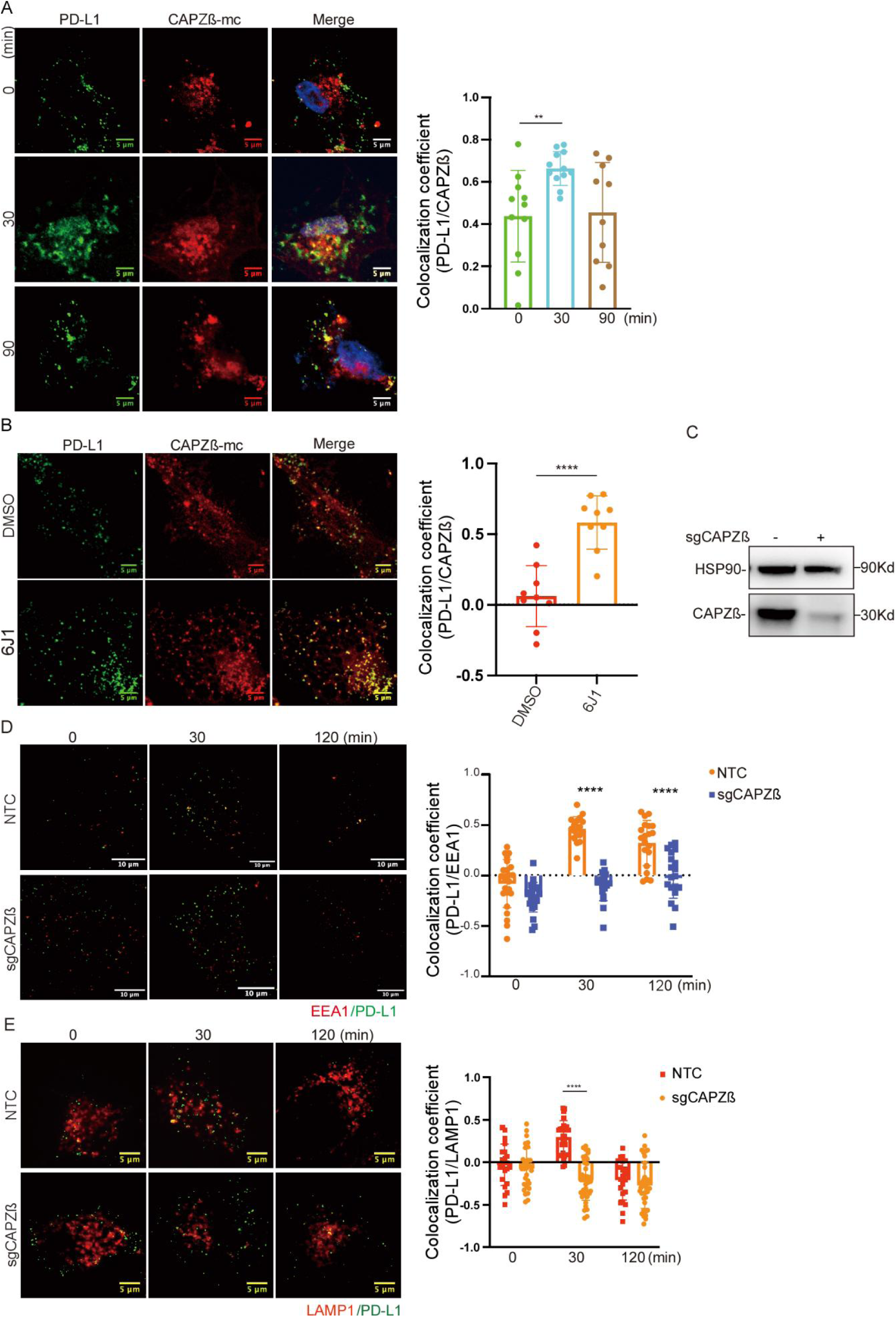
CAPZ is required for endosomal trafficking of internalized PD-L1. (A) HeLa cells expressing CAPZβ-mCherry (CAPZβ-mc) were labeled for cell surface PD-L1 and chased at 37 °C for the indicated times. Representative confocal images and quantification of colocalization of internalized PD-L1 with CAPZβ-positive structures are shown. Scale bars, 5 μm. (B) HeLa cells expressing CAPZβ-mCherry were treated with DMSO or 6J1, followed by analysis of PD-L1 and CAPZβ localization. Representative images and quantification of the colocalization of PD-L1 with CAPZβ-positive structures are shown. Scale bars, 5 μm. (C) Immunoblot analysis of CAPZβ expression in control and CAPZβ-deficient HeLa cells. HSP90 was used as a loading control. (D) Control (NTC) and CAPZβ-deficient HeLa cells were labeled for cell surface PD-L1 and chased for the indicated times, followed by immunostaining for PD-L1 and EEA1. Representative images and quantification of the colocalization of internalized PD-L1 with the early endosomal marker EEA1 are shown. Scale bars, 10 μm. (E) Control (NTC) and CAPZβ-deficient HeLa cells were labeled for cell surface PD-L1 and chased for the indicated times, followed by immunostaining for PD-L1 and LAMP1. Representative images and quantification of the colocalization of internalized PD-L1 with the late endosomal/lysosomal marker LAMP1 are shown. Scale bars, 5 μm. Data are presented as mean ± SEM. Statistical significance was analyzed as indicated in the figure. **p < 0.01; ****p < 0.0001.

To assess the functional contribution of CAPZ, we depleted CAPZβ and first confirmed efficient knockdown by immunoblotting (**Fig. 4C**). Because CAPZ is a key actin-capping protein, we next examined F-actin organization to verify functional disruption. CAPZβ depletion caused a marked increase in actin filament density and the accumulation of aberrant F-actin structures relative to control cells (**Fig. S4),** and quantitative analysis confirmed significantly elevated actin intensity in CAPZβ-deficient cells. These results verify effective disruption of CAPZ function.

We then asked whether CAPZ is required for PD-L1 trafficking through the endosomal pathway. In control cells, internalized PD-L1 robustly colocalized with EEA1-positive early endosomes at 30 min and redistributed over time (**Fig. 4D**). In contrast, CAPZ-deficient cells exhibited markedly reduced PD-L1–EEA1 colocalization at both 30 and 120 min, as reflected by significantly decreased colocalization coefficients. These findings indicate that CAPZ is required for efficient delivery of PD-L1 to early endosomes. Consistent with impaired early endosomal entry, CAPZ depletion also disrupted PD-L1 progression to late endosomal compartments. Whereas control cells displayed expected time-dependent colocalization between PD-L1 and LAMP1-positive structures, CAPZ-deficient cells showed significantly reduced PD-L1–LAMP1 overlap (**Fig. 4E**). Together, these results demonstrate that CAPZ is required for efficient progression of internalized PD-L1 through both early and late endosomal compartments.

Thus, in contrast to autophagy disruption (**Figs. 1–3**), which did not affect PD-L1 trafficking, loss of CAPZ directly impairs PD-L1 delivery to both early and late endosomal compartments. These findings identify CAPZ as a critical determinant of PD-L1 endosomal trafficking.

### CAPZ regulates the exosomal trafficking and secretion of PD-L1

Having established that CAPZ is required for efficient endosomal trafficking of PD-L1 (**Fig. 4**), we next investigated whether CAPZ also participates in exosomal secretion. To determine whether CAPZ is associated with exosome production, we first analyzed exosomal fractions. CAPZβ was detected in purified exosomes isolated from HeLa cells (**Fig. S5**), together with established exosomal markers. These data indicate that CAPZ is present within extracellular vesicles and suggest a potential role in exosome biogenesis or cargo sorting.

We next examined whether CAPZ associates with MVB compartments. CAPZβ-GFP exhibited clear colocalization with CD63-positive vesicles (**Fig. 5A**), indicating that CAPZ localizes to MVB structures involved in exosome biogenesis. This spatial association suggested a potential role for CAPZ in exosomal PD-L1 trafficking. Consistent with impaired endosomal progression observed upon CAPZ depletion (**Fig. 4**), CAPZβ knockdown significantly reduced colocalization between internalized PD-L1 and CD63 at early time points (**Fig. 5B**), indicating defective delivery of PD-L1 to MVB compartments. Quantitative analysis revealed a marked decrease in PD-L1–CD63 correlation coefficients in CAPZ-deficient cells relative to controls. We then examined whether CAPZ regulates exosome composition. Immunoblot analysis of purified exosomes showed that CAPZ depletion markedly reduced the levels of PD-L1 in exosomal fractions and was accompanied by decreased levels of the exosomal markers ALIX, CD63, and CD9 (**Fig. 5C**). These findings indicate that CAPZ is required for efficient exosomal trafficking and/or secretion of PD-L1.

**Figure 5.**
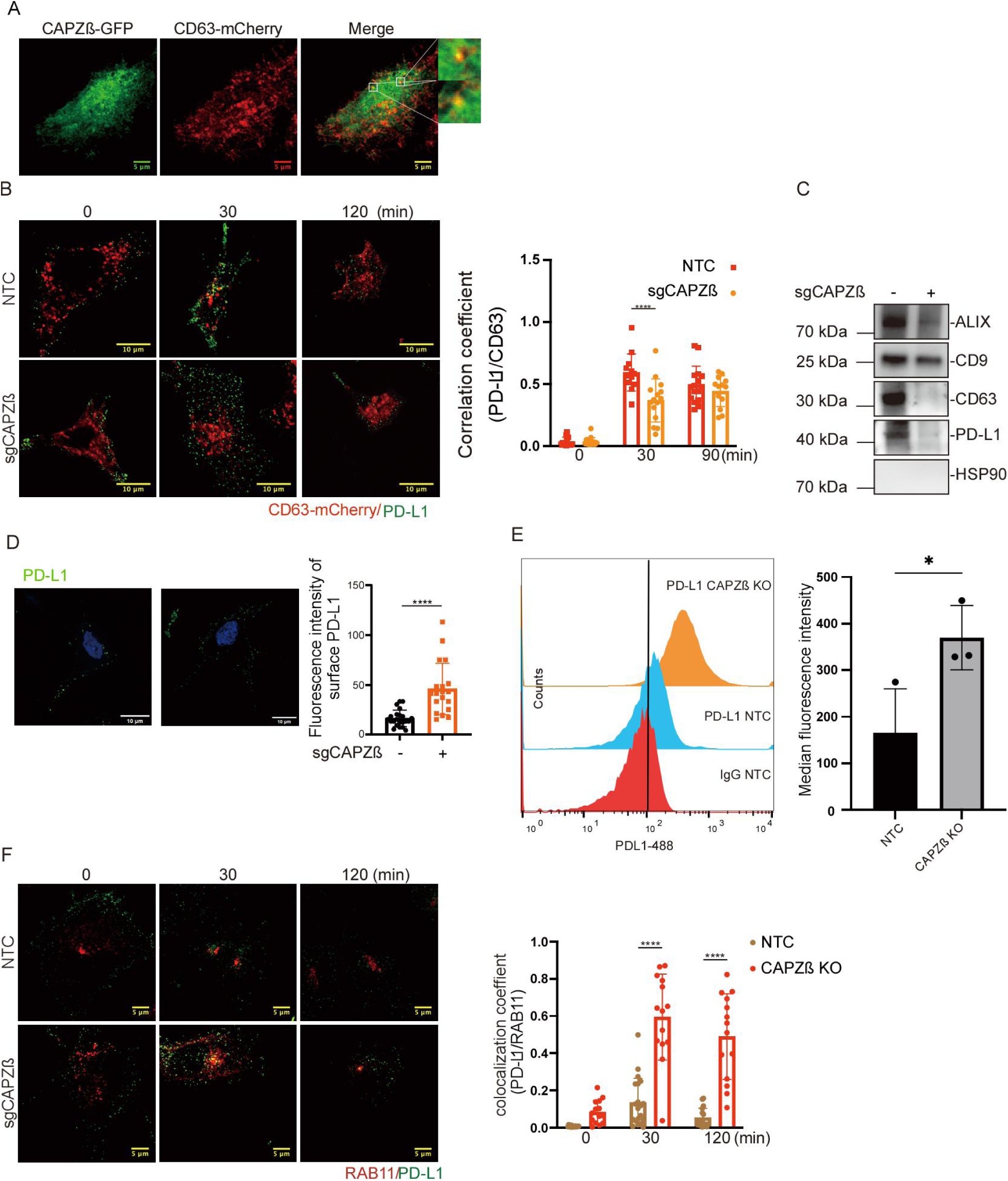
CAPZ promotes incorporation of PD-L1 into CD63-positive compartments and restrains PD-L1 recycling to the plasma membrane. (A) HeLa cells coexpressing CAPZβ-GFP and CD63-mCherry were analyzed by confocal microscopy. Representative images show partial overlap of CAPZβ-positive structures with CD63-positive compartments. Insets show enlarged views of the boxed regions. Scale bars, 5 μm. (B) Control (NTC) and CAPZβ-deficient HeLa cells were labeled for cell surface PD-L1 and chased at 37 °C for the indicated times, followed by analysis of PD-L1 colocalization with CD63-mCherry. Representative images and quantification show the colocalization of internalized PD-L1 with CD63-positive compartments. Scale bars, 10 μm. (C) Exosomes isolated from conditioned medium of control and CAPZβ-deficient HeLa cells were analyzed by immunoblotting for ALIX, CD9, CD63, and PD-L1. HSP90 was used as a negative control. (D) Cell surface PD-L1 in control and CAPZβ-deficient HeLa cells was analyzed by immunofluorescence staining under nonpermeabilizing conditions. Representative images and quantification of cell-surface PD-L1 fluorescence intensity are shown. Scale bars, 10 μm. (E) Cell surface PD-L1 in control and CAPZβ-deficient HeLa cells was analyzed by flow cytometry. Representative histograms and quantification of the surface PD-L1 abundance are shown. IgG staining was used as a negative control. (F) Control (NTC) and CAPZβ-deficient HeLa cells were labeled for cell surface PD-L1 and chased for the indicated times, followed by immunostaining for PD-L1 and RAB11. Representative images and quantification the colocalization of internalized PD-L1 with the recycling endosome marker RAB11 are shown. Scale bars, 5 μm. Data are presented as mean ± SEM. Statistical significance was analyzed as indicated in the figure. *p < 0.05; ****p < 0.0001.

Consistent with reduced exosomal PD-L1 secretion, CAPZ-deficient cells exhibited increased surface PD-L1 abundance. Immunofluorescence analysis revealed significantly elevated surface PD-L1 intensity in CAPZ-depleted cells (**Fig. 5D**), and flow cytometry confirmed a robust increase in cell-surface PD-L1 levels (**Fig. 5E**). These findings suggest that impaired endosomal sorting and exosomal export lead to retention of PD-L1 at the plasma membrane. To further define the trafficking defect, we examined PD-L1 localization relative to RAB11, a marker of recycling endosomes^55^. CAPZ depletion markedly increased PD-L1 colocalization with RAB11 at 30 and 120 min (**Fig. 5F**), indicating enhanced recycling of PD-L1 back to the plasma membrane rather than delivery to MVB/exosomal compartments.

Collectively, these findings establish CAPZ as a critical regulator of exosomal PD-L1 secretion. Unlike canonical autophagy components (Figs. 1–3), which are dispensable for PD-L1 trafficking, CAPZ is required for efficient sorting of PD-L1 into MVB compartments and its subsequent secretion via exosomes.

## DISCUSSION

In this study, we show that the endosomal and exosomal trafficking of cell surface PD-L1 is independent of the canonical autophagy machinery and is instead regulated by CAPZ-dependent endosomal dynamics. Although internalized PD-L1 exhibited substantial colocalization with LC3-positive membranes, particularly under conditions of impaired autophagic flux, genetic disruption of core autophagy components, including LC3B, ATG4B, ATG5, and ATG7, did not impair PD-L1 trafficking to early endosomes, late endosomal/multivesicular body compartments, or exosomes (**Figs. 1–3**). These findings resolve the apparent paradox between the physical association of PD-L1 with LC3-positive vesicles and the lack of functional dependence on canonical autophagy. Specifically, the increased PD-L1–LC3 colocalization observed after pharmacological blockade of autophagic flux most likely reflects vesicular accumulation or compartmental convergence rather than autophagy-dependent sorting. Thus, the presence of LC3 on PD-L1-containing membranes does not indicate that PD-L1 trafficking is mediated by canonical autophagy. Instead, our results support a model in which membrane-derived PD-L1 traffics through endosomal and exosomal pathways independently of canonical autophagy and is functionally controlled by CAPZ-dependent endosomal regulation.

These findings shift the mechanistic framework for PD-L1 trafficking. Rather than being directed by canonical autophagy, PD-L1 sorting is controlled by CAPZ-dependent endosomal dynamics. Consistent with our previous work demonstrating that CAPZ regulates Rab-mediated endosomal maturation and cargo trafficking ^40,41,54^, we show here that CAPZ is required for efficient delivery of internalized PD-L1 to early endosomes and multivesicular bodies. Loss of CAPZ disrupts this routing, reduces PD-L1 incorporation into exosomes, and redirects PD-L1 toward recycling pathways, resulting in increased surface PD-L1 abundance (**Figs. 4-5**). These data position CAPZ as a critical determinant of the sorting decision that governs PD-L1 membrane fate.

Our results also help reconcile previous observations linking autophagy to PD-L1 regulation. Increased PD-L1 expression has been reported in association with Beclin-1 deficiency in pre-B cell leukemia/lymphoma^56^, and several small molecules, including the Sigma-1 inhibitor IPAG and verteporfin, promote PD-L1 degradation through autophagy activation ^57,58^. Conversely, inhibition of autophagy has been reported to increase PD-L1 levels in gastric cancer and osteosarcoma cells^25,59^. Our findings indicate that canonical autophagy does not control the trafficking of surface-derived PD-L1. Instead, autophagy may regulate distinct intracellular PD-L1 pools, such as newly synthesized or misfolded proteins, without influencing the trafficking itinerary of endocytosed PD-L1. Consistent with this interpretation, depletion of ATG4 or ATG7 did not significantly alter basal PD-L1 abundance in our system. Moreover, ionomycin-induced Ca²⁺ elevation markedly reduced total PD-L1 levels, likely reflecting stimulated extracellular vesicle release, but this reduction was not prevented by autophagy gene disruption, further supporting the autophagy-independent nature of PD-L1 membrane trafficking (**Figs. S3C-S3D**).

Although canonical autophagy is dispensable for PD-L1 trafficking, noncanonical ATG8 conjugation pathways may still contribute to this process. In conjugation of ATG8 to single membranes (CASM) and related pathways such as LC3-associated phagocytosis (LAP) and LC3-associated endocytosis (LANDO), ATG8 family proteins are lipidated and recruited to single membranes involved in endocytic or phagocytic processes. These pathways use a ubiquitin-like conjugation system that overlaps substantially with that of canonical autophagy ^60,61^. In mammals, the ATG8 family includes the LC3 subfamily (LC3A, LC3B, LC3C) and the GABARAP subfamily (GABARAP, GABARAPL1, and GABARAPL2)^62^. In our study, LC3B localized to PD-L1–containing early endosomes and multivesicular bodies, particularly in cells treated with triazine compounds (**Figs. 1-2**). However, LC3B depletion did not affect PD-L1 trafficking, suggesting that other ATG8 family members may be recruited to PD-L1–containing endosomal compartments. Such recruitment could influence sorting decisions that determine whether PD-L1 is recycled to the plasma membrane, delivered to lysosomes, or incorporated into exosomes. Future studies will therefore be required to determine whether other ATG8 family members participate in PD-L1 trafficking through redundant or noncanonical mechanisms.

In summary, our findings redefine the regulatory logic of PD-L1 membrane trafficking. Canonical autophagy does not determine the trafficking fate of endocytosed PD-L1. Instead, CAPZ-dependent endosomal regulation governs the sorting decisions that direct PD-L1 toward recycling or exosomal secretion. This mechanistic framework highlights endosomal trafficking networks, rather than autophagy, as central regulators of immune checkpoint distribution and suggests new opportunities for modulating PD-L1–mediated immune responses.

## EXPERIMENTAL PROCEDURES

### Cell Culture

HeLa cells (ATCC) and 4T1 mouse mammary carcinoma cells (kindly provided by Dr. Minh Le) were maintained in Dulbecco’s Modified Eagle Medium (DMEM; Invitrogen, 12800-017) supplemented with 10% fetal bovine serum (FBS; Invitrogen, 10270-106) and 100 U/ml penicillin–streptomycin (Invitrogen, 15140-122). Cells were cultured at 37 °C in a humidified incubator containing 5% CO₂. Cells were routinely tested for mycoplasma contamination and used within 10–15 passages after thawing. For experiments involving extracellular vesicle (EV) collection, cells were cultured in medium supplemented with EV-depleted FBS prepared by ultracentrifugation at 100,000 × g for 16 h.

### Reagents and Antibodies

The following primary antibodies were used in this study: anti-Rab5 (#3547), anti-Rab27a (#69295), anti-Rab11 (#5589), anti-PD-L1 (#13684), anti-ATG5 (#2630), anti-ATG4B (#5299), and anti-ATG7 (#8558), all from Cell Signaling Technology. Antibodies against TSG101 (14497-1-AP), CD63 (25682-1-AP), Alix (12422-1-AP), Calnexin (10427-2-AP), GAPDH (60004-1-Ig), and β-actin (66009-1-Ig) were purchased from Proteintech. Fluorescently labeled anti-PD-L1 antibodies were obtained from Thermo Fisher Scientific (MIH1, 16-5983-38; MIH5, 16-5982-38). Bafilomycin A1 was purchased from Selleck Chemicals and dissolved in DMSO as a stock solution.

### Immunofluorescence Microscopy

For labeling of cell surface PD-L1 in live cells, cells were washed with cold phosphate-buffered saline (PBS) and incubated with anti-PD-L1 antibody diluted in PBS containing 0.5% bovine serum albumin (BSA) for 1 h on ice to prevent endocytosis. Cells were washed three times with ice-cold PBS containing 0.5% BSA and incubated with fluorophore-conjugated secondary antibodies for 1 h on ice. After labeling, cells were washed with PBS prior to imaging or further processing.

For intracellular staining, cells grown on coverslips were fixed with 4% paraformaldehyde (PFA) in PBS for 15 min at room temperature, followed by permeabilization with 0.3% Triton X-100 in PBS for 15 min. Cells were then blocked with 5% BSA in PBS for 1 h at room temperature and incubated with primary antibodies overnight at 4 °C. After three washes with PBS, cells were incubated with appropriate fluorescence-conjugated secondary antibodies for 1 h at room temperature. Nuclei were stained with DAPI for 5 min. Coverslips were mounted using ProLong Diamond Antifade Mountant (Thermo Fisher Scientific, P36970).

Images were acquired using either a Zeiss LSM880 confocal microscope or a Nikon A1HD25 high-speed confocal microscope equipped with a large-field-of-view detector. Image acquisition parameters were kept constant within each experiment. Image processing and analysis were performed using Zeiss ZEN software or Nikon NIS-Elements AR software.

### Flow Cytometry

Cell surface PD-L1 expression was quantified by flow cytometry. Live cells were harvested and resuspended in PBS containing 1% BSA. Cells were incubated with anti-PD-L1 antibody for 30 min at 4 °C. After washing twice with PBS, fluorescence signals were analyzed using a Beckman Coulter CytoFLEX flow cytometer. Data were analyzed using FlowJo software (version 10). Dead cells were excluded based on forward and side scatter parameters.

### CRISPR-Cas9 Gene Disruption

Single guide RNAs (sgRNAs) targeting autophagy genes were designed using the Benchling CRISPR design platform. Oligonucleotides encoding sgRNAs were cloned into the lentiCRISPRv2-puro vector. The following sgRNA sequences were used: ATG4B: AGCAAACCGGAGAGTGTCGT; ATG5: AACTTGTTTCACGCTATATC; ATG7: AGAAGAAGCTGAACGAGTAT; LC3B: CGGAGAAGACCTTCAAGCAG; Beclin1: AAAGGAGCTGGCACTAGAGG. Lentiviral particles were produced in HEK293T cells using standard packaging plasmids. Target cells were infected with lentivirus and selected with puromycin (2 µg/ml) for 5–7 days to generate stable knockout populations. Gene disruption efficiency was verified by immunoblot analysis.

### Extracellular Vesicle Isolation

Extracellular vesicles were isolated by differential ultracentrifugation. Cells were cultured in EV-depleted medium and treated with DMSO (vehicle control) or 6J1 (1 µM) for 48 h. Conditioned medium was collected and subjected to sequential centrifugation to remove cells and debris: 1,000 × g for 10 min, 2,000 × g for 15 min, and 10,000 × g for 15 min at 4 °C. The cleared supernatant was then ultracentrifuged at 110,000 × g for 90 min using a Beckman ultracentrifuge. The pellet containing extracellular vesicles was washed in 10 ml cold PBS and subjected to a second ultracentrifugation at 110,000 × g for 90 min. The final EV pellet was resuspended in PBS and used for downstream analyses including immunoblotting and nanoparticle tracking analysis.

### Statistical Analysis

Data are presented as mean ± standard error of the mean (SEM). Statistical comparisons between two groups were performed using an unpaired two-tailed Student’s t test. Multiple comparisons were analyzed using one-way ANOVA followed by appropriate post hoc tests. Statistical significance was defined as *P* < 0.05.

### Data Availability

All data supporting the findings of this study are included within the manuscript and supplementary materials. Plasmids generated in this study are available from the corresponding author upon reasonable request.

## ACKNOWLEDGMENTS

We thank members of Yue lab for their advice on preparing this manuscript. This work was supported by NSFC (32570826 and 82301945), Jiangsu Provincial Natural Science Foundation (BK20251832), Kunshan Shuang Chuang Grant (kssc202302073), Suzhou Innovation and Entrepreneurship Leading Talent Program (ZXL2024337), and research grants from Shenzhen Science and Technology Innovation Committee (SGDX20201103093201010 and JCYJ20210324134007020).

---

**Figure S1.**
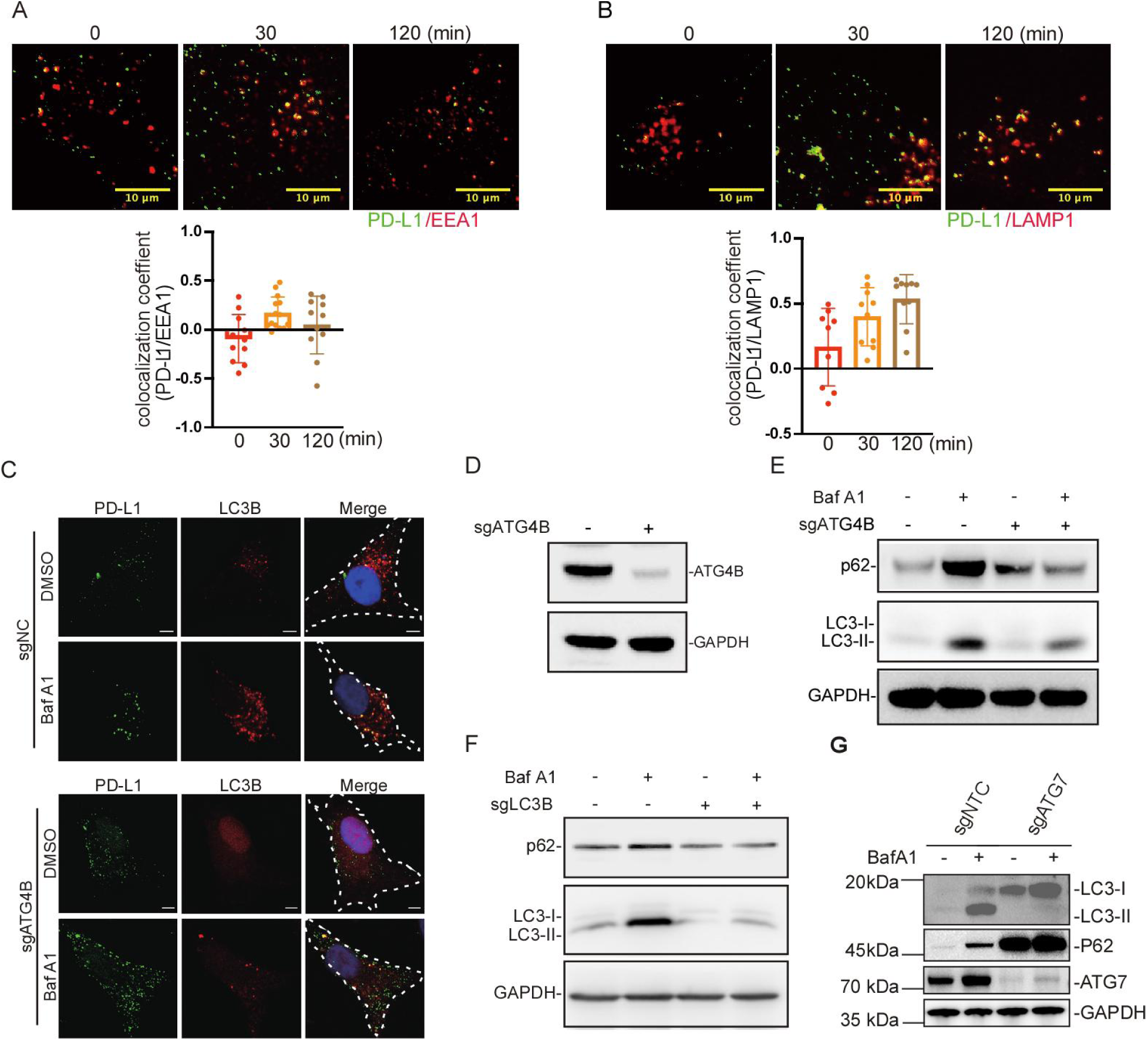
Validation of PD-L1 endosomal trafficking and autophagy perturbation. (A) Colocalization of internalized PD-L1 with EEA1 at indicated time points. (B) Colocalization of internalized PD-L1 with LAMP1. (C) Colocalization of PD-L1 with LC3B under basal conditions or following Bafilomycin A1 treatment, and in ATG4B-deficient cells. (D–E) Immunoblot validation of ATG4B knockdown and assessment of autophagic flux markers (LC3, p62). (F) Immunoblot validation of LC3B knockdown and assessment of autophagic flux markers (LC3, p62). (G) Immunoblot validation of ATG7 depletion and assessment of autophagic flux markers (LC3, p62).

**Figure S2.**
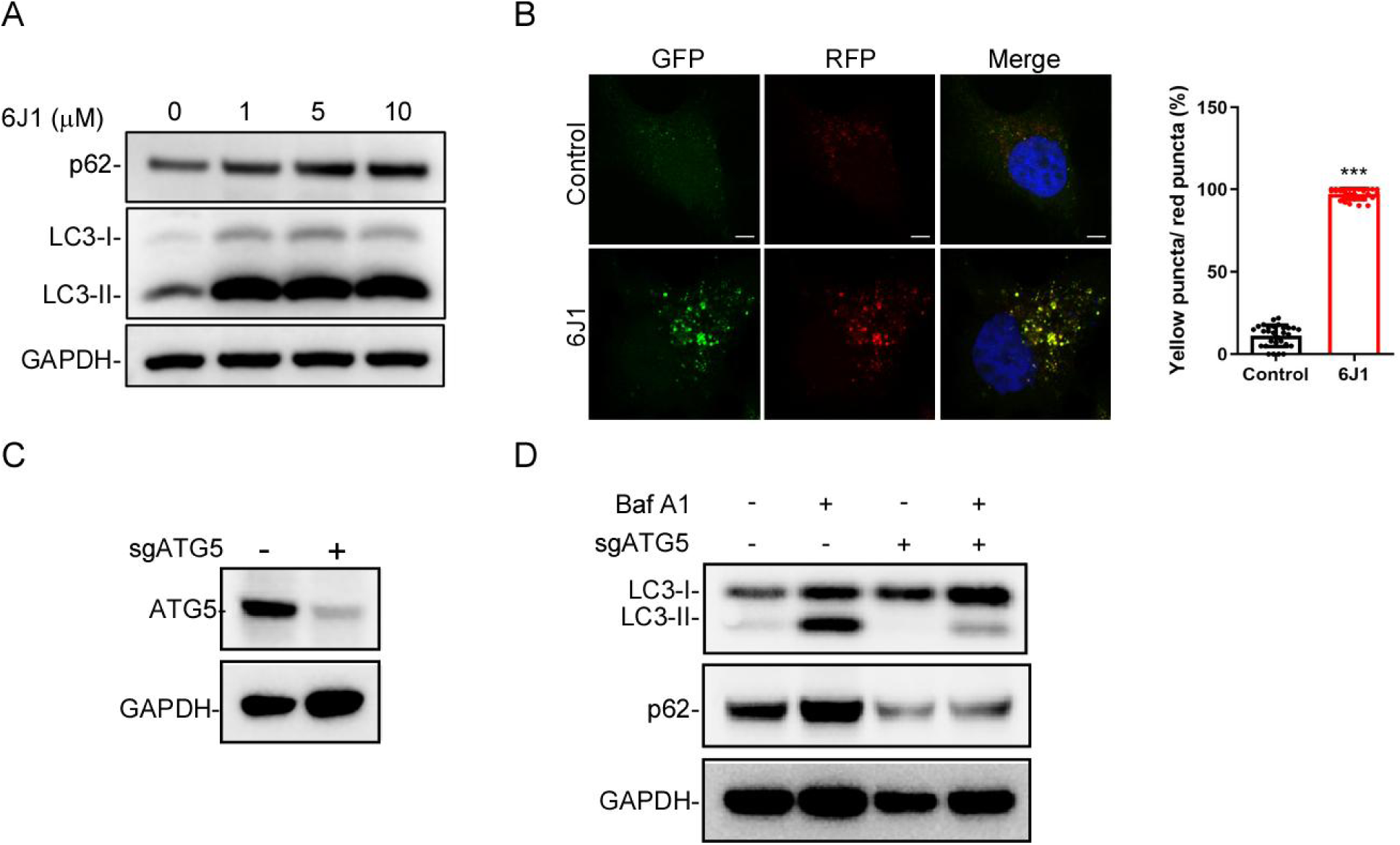
6J1 is a late-stage autophagy inhibitor, and ATG5 knockdown abolished autophagic flux. **(A)** HeLa cells were treated with or without 6J1 for 6 h, after which they were collected for western blot analysis of LC3 and p62. **(B)** GFP-RFP-LC3 expressing HeLa cells were plated on coverslips in 24-well plates and treated with or without 6J1 (1 μM) for 6 h, after which the cells were fixed and imaged. The scale bar is 5 μm. MCC of LC3-II green puncta with LC3-II red puncta were quantified. (**C**) The knockdown efficiency of ATG5 in HeLa cells. (**D**) Control or ATG5 knockdown HeLa cells were treated with or without Baf-A1 (10 nM) for 12 h, followed by immunoblot analysis against LC3B, SQSTM1 (p62), and GAPDH.

**Figure S3.**
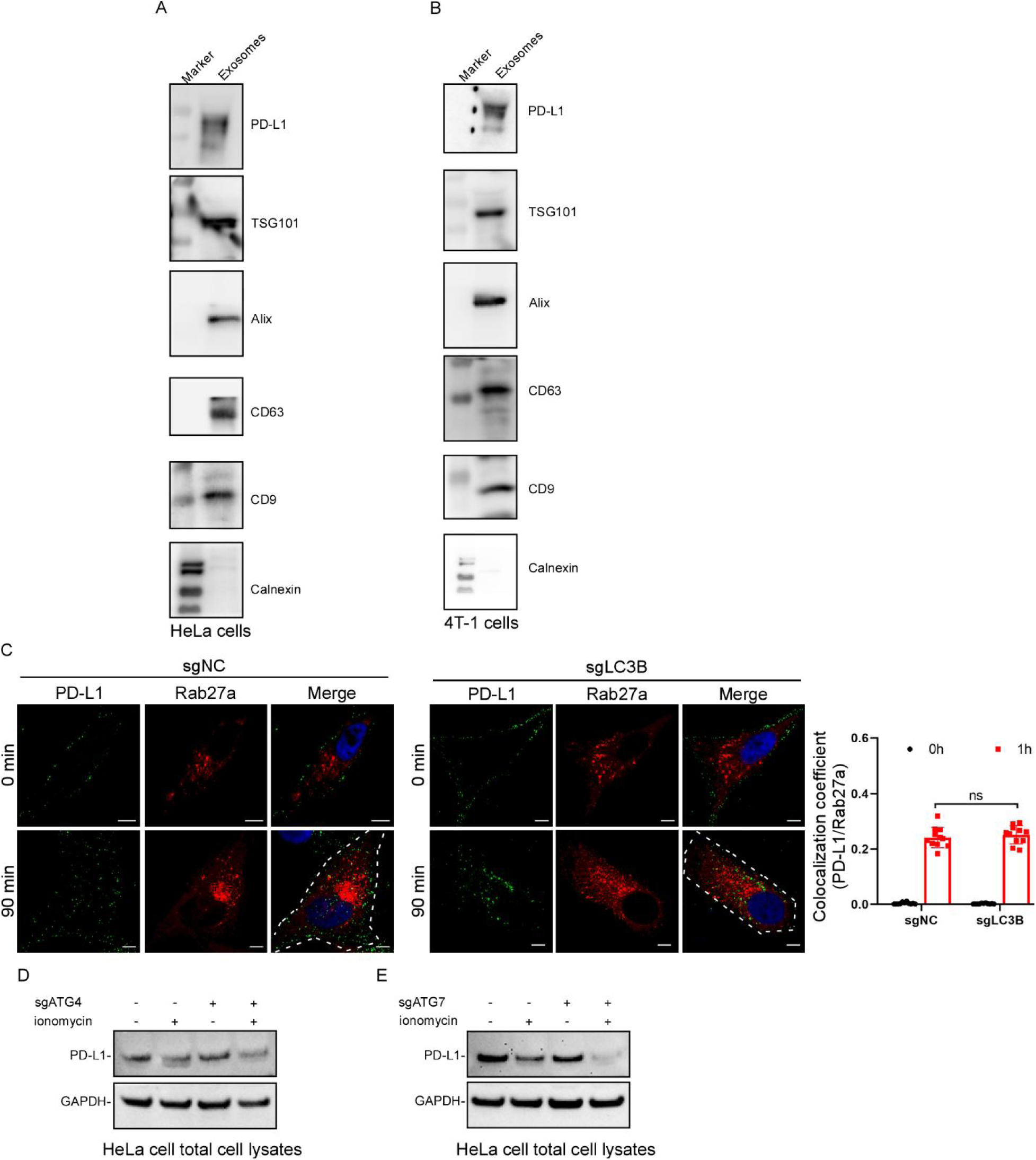
**Canonical autophagy is not required for exosomal trafficking and secretion of membrane PD-L1. (**A, B) EVs isolated from HeLa (A) or 4T1 (B) cells were analyzed by immunoblotting for PD-L1 and the exosomal markers TSG101, ALIX, CD63, and CD9. Calnexin was included as a negative control for endoplasmic reticulum contamination. (C) Control and LC3B-deficient HeLa cells were labeled for cell surface PD-L1 and chased for the indicated times, followed by immunostaining for PD-L1 and Rab27A. Representative confocal images and quantification of the internalized PD-L1 colocalized with Rab27A are shown. Scale bars, 10 μm. (D, E) Control and ATG4B-deficient (D) or ATG7-deficient (E) HeLa cells were treated with ionomycin for the indicated conditions, and total cell lysates were analyzed by immunoblotting for PD-L1. GAPDH was used as a loading control.

**Figure S4.**
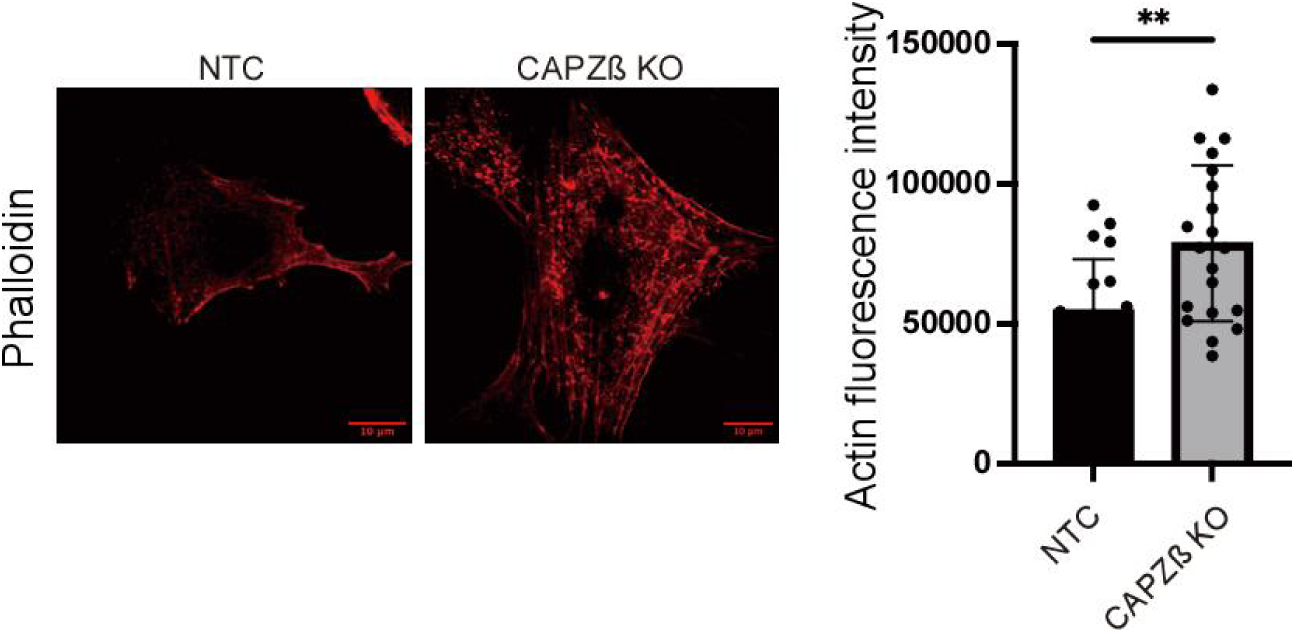
CAPZ depletion increases actin filament density. Control (NTC) and CAPZβ-deficient HeLa cells were stained with phalloidin to visualize F-actin. Representative confocal images show increased actin filament density and accumulation of aberrant F-actin structures in CAPZβ-deficient cells compared with control cells. Quantification of integrated actin density confirmed a significant increase in F-actin signal following CAPZβ depletion. Scale bars, 10 μm. Data are presented as mean ± SEM. **p < 0.01.

**Figure S5.**
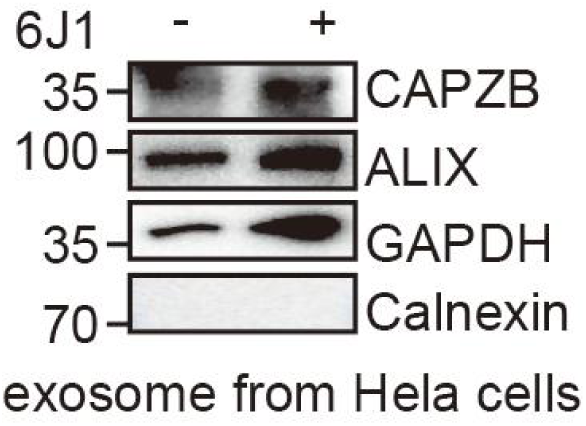
CAPZ is present in exosomal fractions. Immunoblot analysis showing CAPZβ and exosomal markers in purified extracellular vesicles.

## Notes

### Competing Interest Statement

The authors have declared no competing interest.

### Summary of Updates

This revised version updates Supplementary Figure S5 with new experimental data to further support the core findings of the study. No other major changes to the main text, figures, or conclusions have been made.

